## Supplementay_Information_Figures for "Single-cell RNA sequencing on formalin-fixed and paraffin-embedded (FFPE) tissue identified multi-ciliary cells in breast cancer"

**Supplementary information**

**Methods**

Processing of fresh tissue samples

Fresh tissue samples were placed immediately on a glass plate over ice and then cut into small pieces with a diameter of 1 mm. Subsequently, the tissue fragments were fixed overnight for 19 hours and were dissociated following the “Tissue Fixation & Dissociation protocol for Chromium Fixed RNA Profiling” (CG000553 from 10x Genomics, Pleasanton, California, USA). The dissociation step was performed automatically using the gentleMACS Octo Dissociator with heaters (Miltenyi) following the program for approximately 35 minutes specified in the protocol. Finally, the cells were counted by staining cells with propidium iodide and using an automated fluorescence cell counter (RWD C100). Single-cell suspensions were then frozen as described in the protocol for long-term storage at -80ºC for 1 month. Subsequently, the protocol was continued with the “Chromium Fixed RNA Profiling Reagent Kits for Singleplexed Samples” (CG000477 from 10X Genomics), following the manufacturer’s protocol without any adjustments. (Figure 1a)

Processing of FFPE tissue samples

FFPE tissue samples were processed following the protocol “Isolation of Cells from FFPE Tissue Sections for Chromium Fixed RNA Profiling” (CG000632 from 10X Genomics). Tissue dissociation was carried out using the gentleMACS Octo Dissociator with heaters (Miltenyi), utilizing the preinstalled program 37C_FFPE_1 for approximately 48 minutes as specified in the protocol. Once the dissociation protocol was completed, the cells were counted and frozen using the same procedure as for cells derived from fresh samples. In this case, the cells were kept at -80ºC for only one week before commencing the “Chromium Fixed RNA Profiling Reagent Kits for Singleplexed Samples protocol” (CG000477 from 10X Genomics). (Figure 1a)

10X Library preparation and sequencing

The samples were thawed and recounted. Single-cell library preparation was conducted following the manufacturer's protocol for the Chromium Fixed RNA Profiling Reagent Kits for Singleplexed Samples (CG000477 from 10X Genomics). The cells were loaded onto the Chromium X/iX to generate single-cell gel beads in emulsions (GEMs). Subsequently, the libraries were sequenced on a NovaSeq 6000 system (Illumina, USA) with an approximate 200 million reads per library. In Supplementary Table 1 and Supplementary Table 2, the number of cells per sample at various stages of the process and a summary of scRNA-seq data obtained per case are detailed.

scRNA-seq data processing

Sequencing reads were aligned against reference transcriptome GRCh38 and unique molecule identifiers (UMIs) were quantified using Cell Ranger, version 7.1.0 (10x Genomics). Subsequent analyses were performed using R (version 4.3.2) and the Seurat package (version 5.0.2). Gene expression data of all samples were merged and filtered for the following quality parameters: >200 and <10,000 genes per cell, >250 and <5,000 UMIs per cell, fraction of mitochondrial reads lower than 20% (Figure S1a,b). After filtering, a total of 46,666 cells remained (Figure S1d). Next, the doublets were analyzed and removed using doubletFinder (version 2.0.4) ^1^, a total of 43,599 cells remained (Figure S1e,f). (Supplementary Table 2)

Gene expression data was log normalized. After performing principal component analysis (PCA), data integration was carried out using Harmony (version 1.2.0) ^2^ to correct batch effects associated with sample type (FFPE or fresh) (max.iter.harmony = 10, lambda = 1). Uniform manifold approximation projection (UMAP) based on the top 15 principal components (PCs) was then used for data visualization and cells were clustered running the “FindNeighbors” and “FindClusters” functions (resolution = 0.15). Main cell types and cell subtypes were manually annotated using canonical marker genes selected from the literature (Figure S2). For a more comprehensive annotation of epithelial, mesenchymal and immune cells, the populations were further extracted and underwent a subanalysis. Cells were reclustered with the “FindClusters” function (dims.use = 1:30, resolution = 0.15) and were visualized in two dimensions using UMAP. The SingleR R package (version 2.4.1) with reference to Monaco (MonacoImmuneData) ^3^ dataset was applied to annotate immune cells. Finally, we used the "FindAllMarkers" function of the Seurat package to distinguish differentially expressed gene (DEGs) in different cell clusters (only.pos =T, min.pct =0.1, logfc. threshold =0.25).

Functional enrichment analysis

Gene Oncology (GO) enrichment analysis to characterize biological processes (BP). Based on genes showing significant positive expression in the cluster of interest, the “enrichGO” function in the clusterprofiler package ^4^ was used to perform functional enrichment analysis. The relevant parameters were set as follows: keyType = “SYMBOL”, pvalueCutoff = 0.01, qvalueCutoff = 0.5, and ont = “BP”.

Inference of copy number variation (CNV) from single-cell RNA-seq

In each tumor, putative copy number events were inferred for each epithelial cell clusters using the R package inferCNV version 1.18.1 ^5^. we used non-malignant cells including immune cells and basal cells as baselines to estimate the CNA of malignant cells. Genes were sorted by their genomic locations on each chromosome. The standard inferCNV algorithm was invoked with infercnv::run() with cutoff set to ‘‘0.1,’’ denoise set to ‘‘TRUE’’ and HMM set to ‘‘TRUE.’’ The default i6 Hidden Markov Model (HMM) was used to predict CNV levels based on a six-state CNV model ranging from complete loss to > 2 copies. The Bayesian Network Latent Mixture Model was used to estimate the posterior probability of each CNV level at each predicted CNV region.

Immunohistochemistry and image analysis

FFPE tissue sections of 3 μm were prepared for IHC. For molecular classification, the two FFPE cases underwent an immunohistochemical study for the expression of estrogen receptors (ER), progesterone receptors (PR), HER2 and Ki67. Evaluation and interpretation of ER, PR, and HER2 expression was performed according to American Society of Clinical Oncology and the College of American Pathologists (ASCO-CAP) guidelines ^6^. In both carcinomas, the expression of E-cadherin was studied to confirm the histological type. In addition, the expression of P63, FOXJ1 was evaluated in both cases and FOXJ1 in the TMA tumors. Furthermore, for the study of immune cell populations, the expression of CD3, CD4, CD8, CD20, CD68 and CD117 was analyzed. (Table of antibodies below).

| **Protein** | **Clon** |  |
| --- | --- | --- |
| ER | 6F11 | Leica Biosystems |
| PR | 16 | Leica Biosystems |
| HER2 | 4B5 | Ventana, Roche Diagnostics |
| E-cadherin | 36b5 | Leica Biosystems |
| P63 | 7 JUL | Leica Biosystems |
| FOXJ1 | EPR21874 | Abcam |
| CD4 | 4B12 | Leica Biosystems |
| CD8 | 4B11 | Leica Biosystems |
| CD3 | LN10 | Leica Biosystems |
| CD20 | L26 | Leica Biosystems |
| CD68 | 514H12 | Leica Biosystems |
| CD117 (c-KIT) | EP10 | Leica Biosystems |

Fluorescent In-Situ Hybridization (FISH)

Chromosomal alterations were evaluated by FISH on FFPE sample slides using the following probes: HER2/17q and MDM4/1p12 dual color Probe Kit (Zytovision GmbH, Bremen, DE). FISH slides were observed with a fluorescence microscope at 100X with immersion oil. A detailed scoring of at least 20 neoplastic cells per sample was performed. Amplification was considered when the tumor cell population had at least twice as many gene signals than centromere signals of the respective chromosome (ratio 2).

**Supplementary Figures**

Supplementary Figure 1. Quality control parameters. (a) Gene expression data, numbers of UMIs and fractions of mitochondrial reads per cell merged before filtering and colored by sample origin. (b) Gene expression data, numbers of UMIs and fractions of mitochondrial reads per cell merged after filtering for the quality parameters: >200 and <10,000 genes per cell, >250 and <5,000 UMIs per cell, fraction of mitochondrial reads lower than 20%. (c) Barcode Rank Plots showing the count of filtered UMIs mapped to each barcode. The color of the graph represents the local density of barcodes that are cell-associated. The plots demonstrated a high quality based on their profiles, all showing a distinctive "cliff and knee" shape. (d) Bar chart showing the number of cells per case after filtering. (e) UMAP plot of 43,599 individual cells from 4 samples generated from the top 15 principal components of all single-cell transcriptomes post-filtering, doublets removing and data integration, colored by singlets and doublets. (f) Bar plot showing singlets and doublets proportions in FFPE and fresh samples. (g) UMAP plot of 43,599 individual cells after doublet removal, colored by tissue origin. (h) Number of genes detected per cell in each cellular population in FFPE and fresh samples.

(h)

(g)

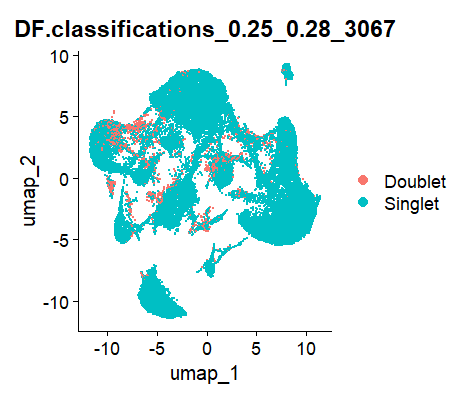

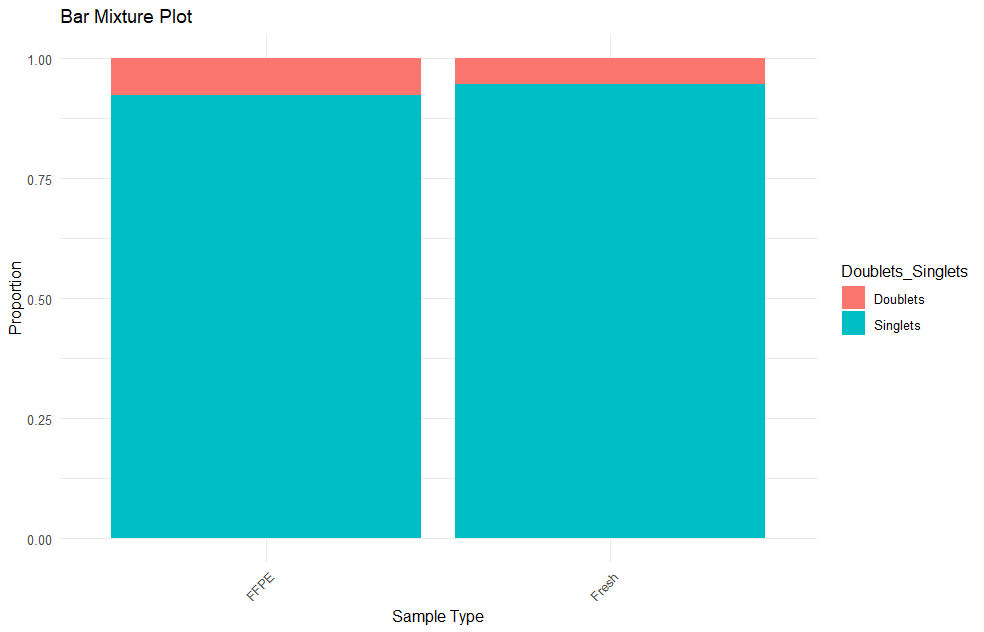

(f)

(e)

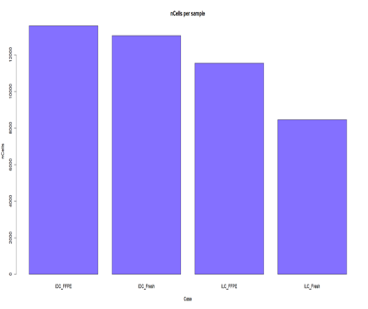

(d)

(a)

(b)

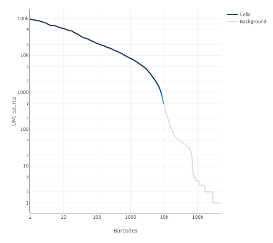

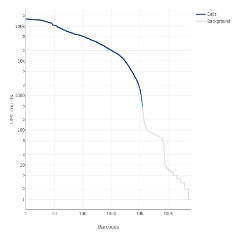

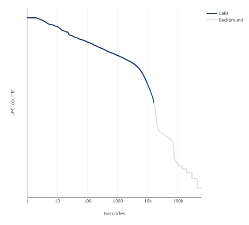

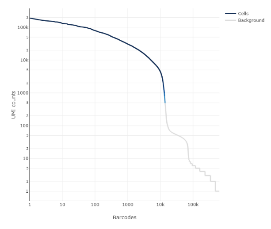

FF ILC

FFPE ILC

FFPE IDC

FF IDC

(c)

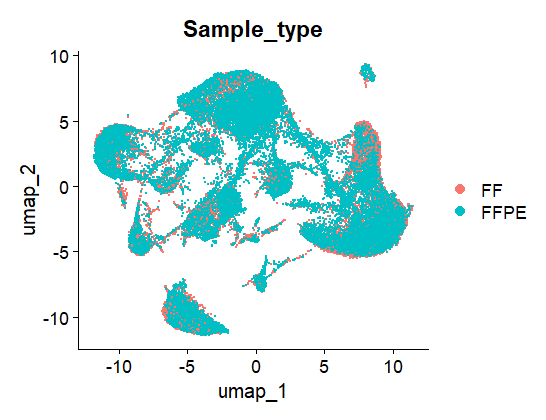

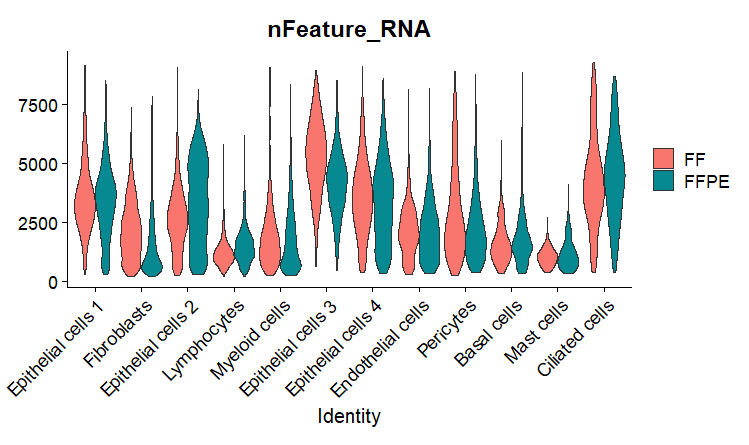

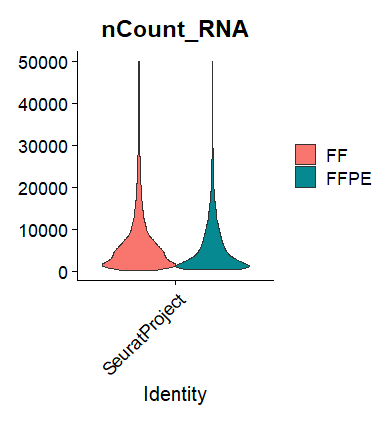

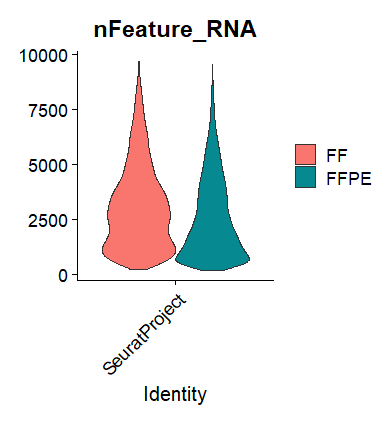

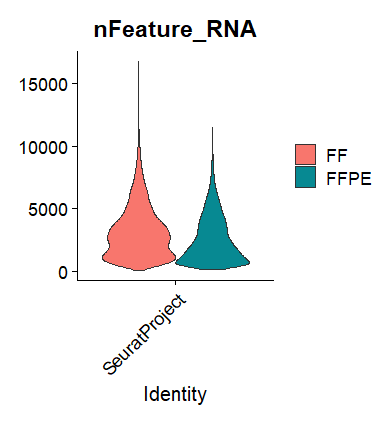

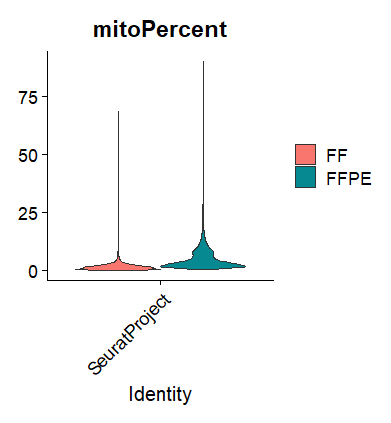

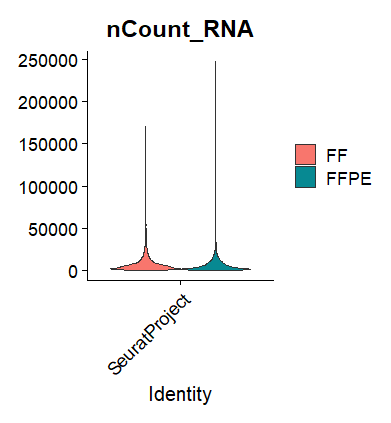

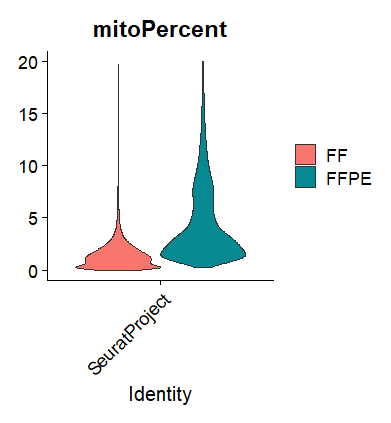

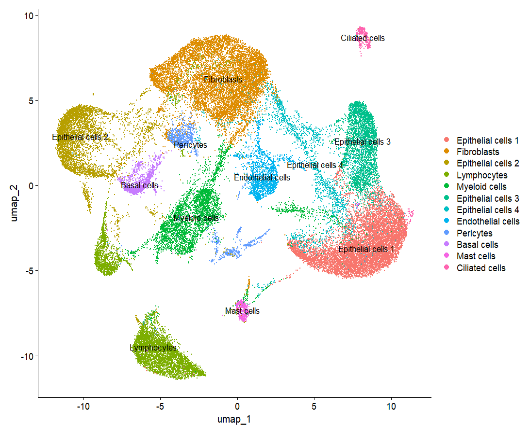

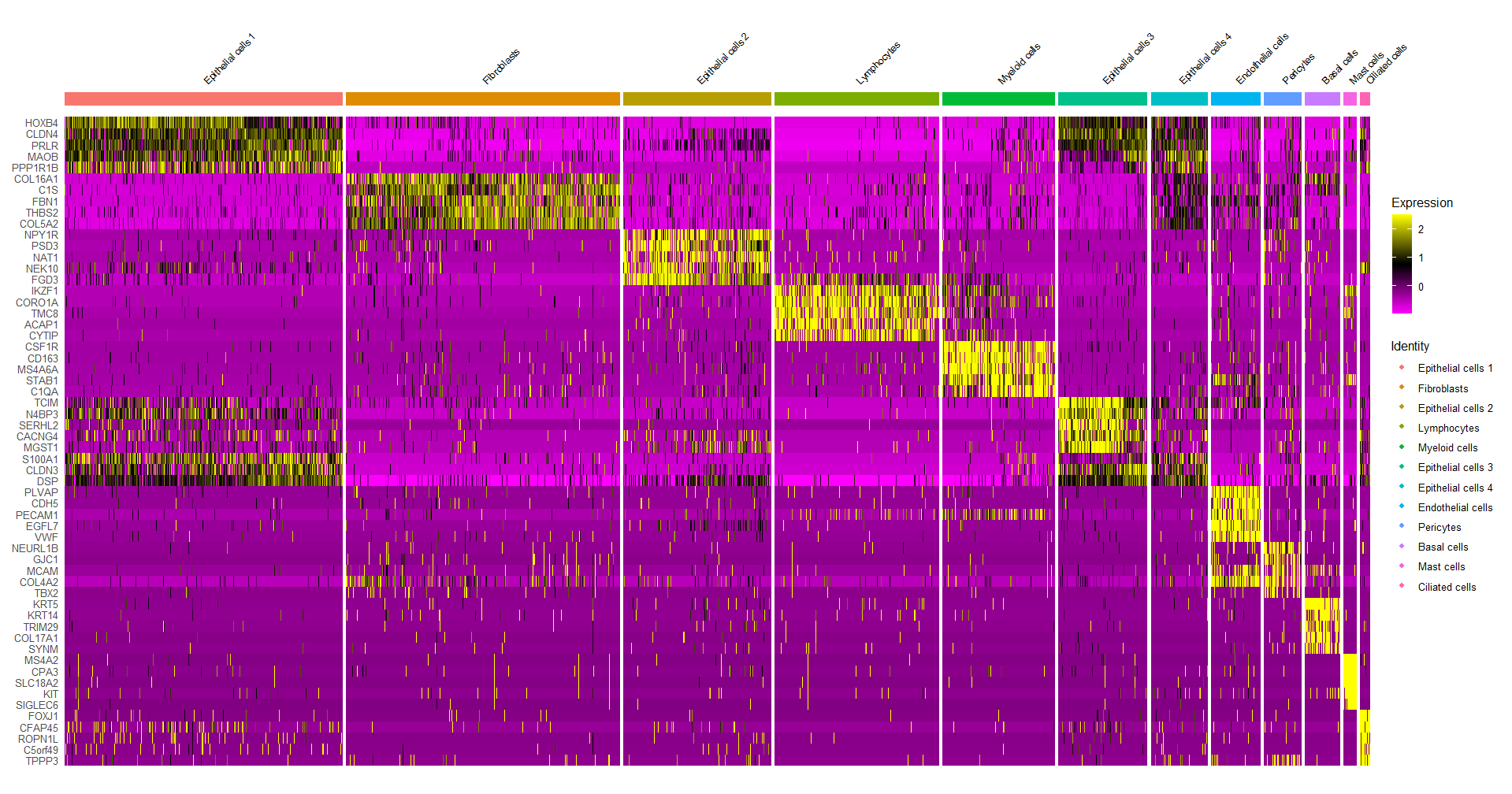

(c)

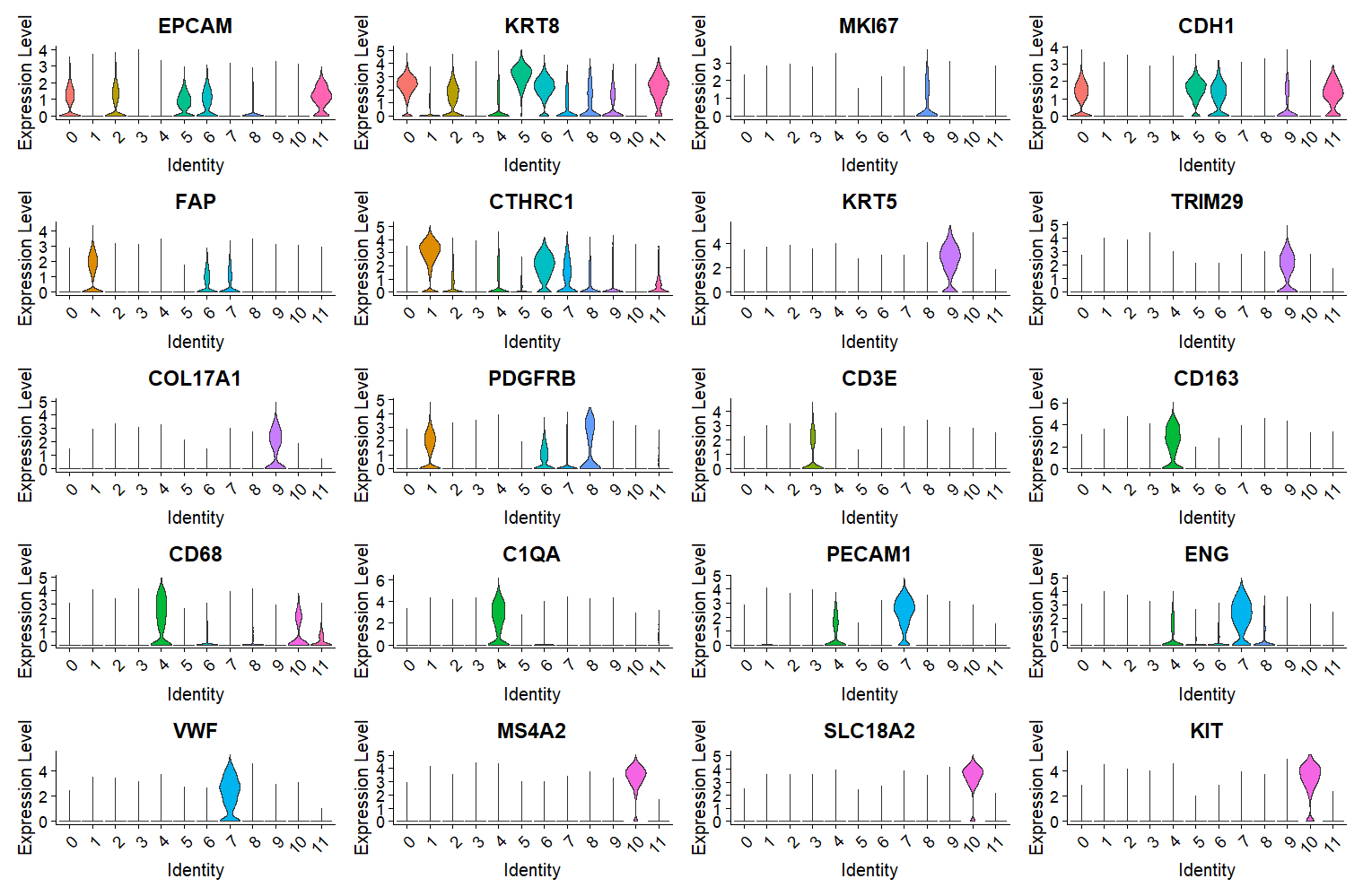

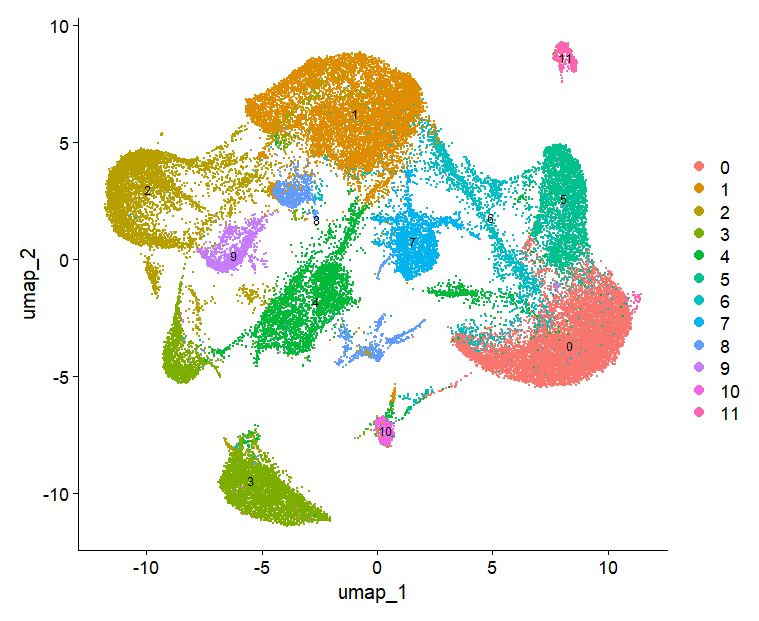

(b)

(a)

(d)

Supplementary Figure 2. scRNAseq annotation. (a) Clustered UMAP plot generated from the top 15 principal components of all single-cell transcriptomes post-filtering and data integration. (b) Markers for cell type annotation. (c) UMAP plot annotated according to the cell markers identified in Figure c. (d) Heatmap representing the top 10 most overexpressed genes (avg_log2FC > 1) in each cell population.

(a)

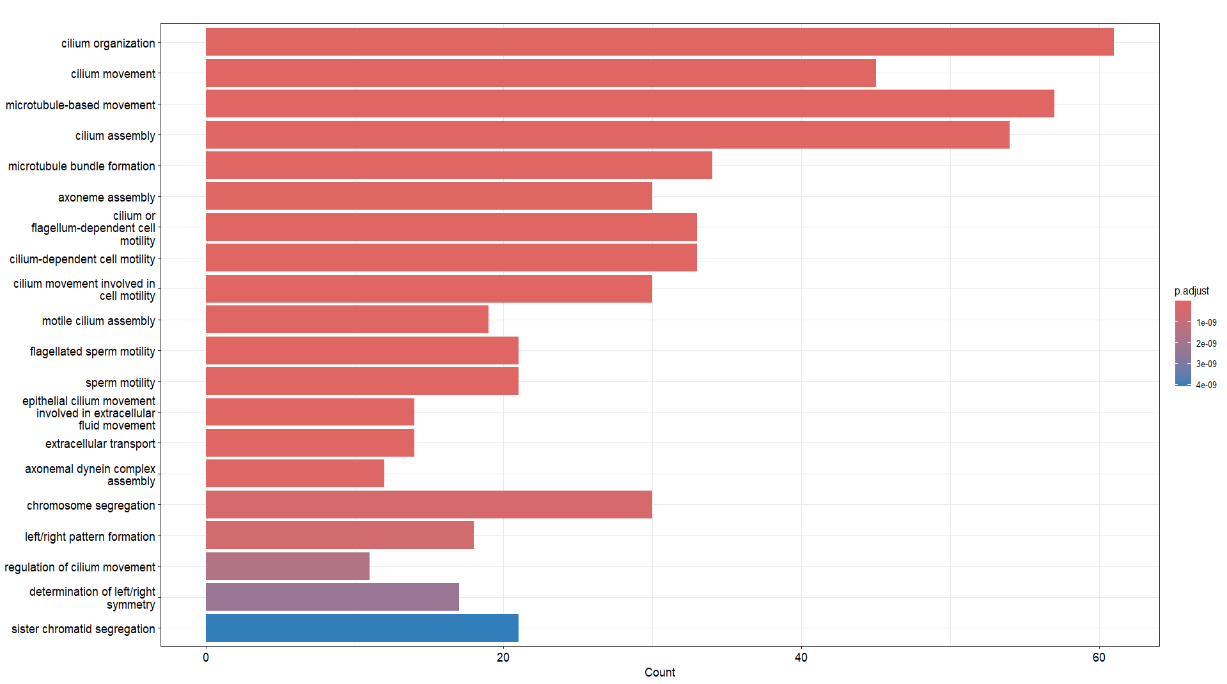

(b)

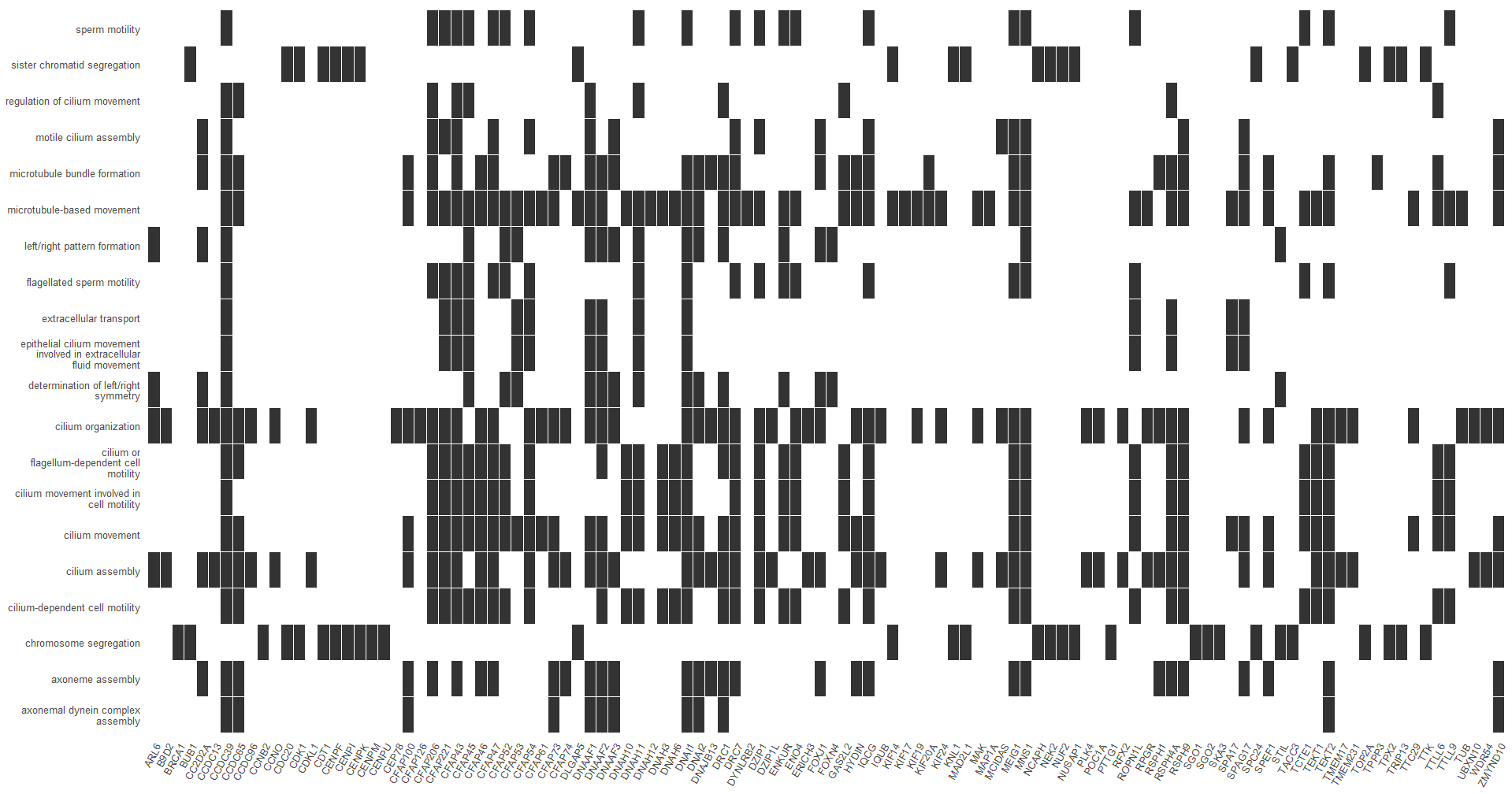

Supplementary Figure 3. GO-enriched biological processes in multiciliated cells. (a) Bar plot displaying Go-enriched pathways significantly upregulated in multiciliated cells. (b) Heatmap showing the GO-enriched biological processes alongside the upregulated genes in multiciliated cells that contribute to these processes.

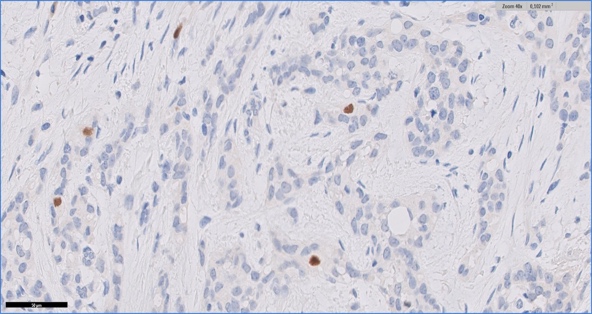

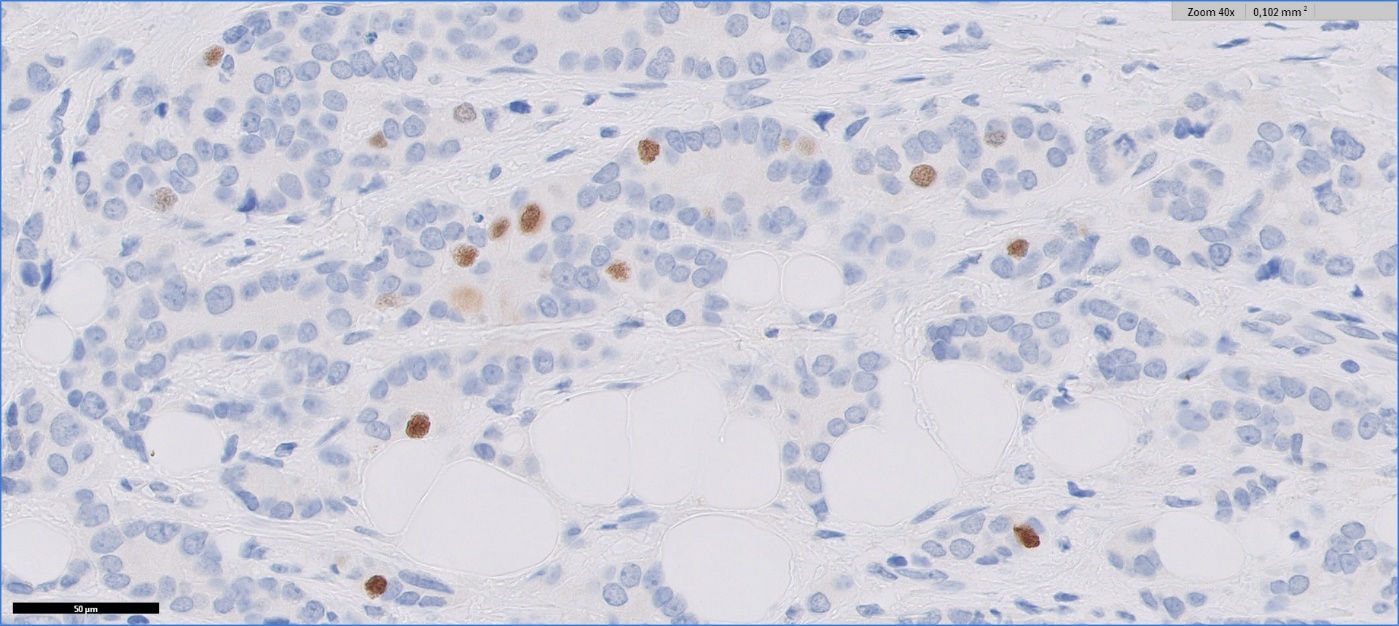

IDC luminal series

(g)

(i)

)

ILC luminal series

(g)

(i)

)

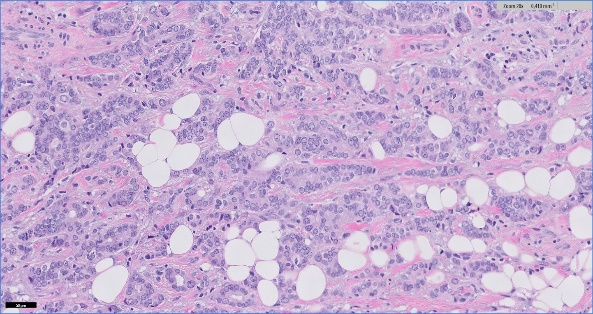

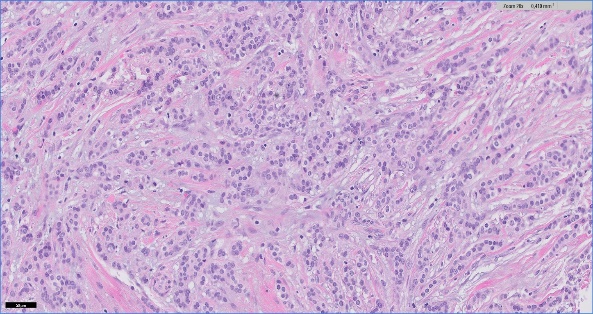

Supplementary Figure 4. H&E staining, and FOXJ1 expression evaluation in one ILC and one IDC of the luminal breast carcinomas series (scale bar 50 μm).
